# Mitochondrial complex I as a diagnostic and therapeutic target in a mouse model of tauopathy

**DOI:** 10.1101/2023.08.13.552232

**Authors:** Jia Hui Wong, Anselm S. Vincent, Shivashankar Khanapur, Tang Jun Rong, Boominathan Ramasamy, Siddesh Hartimath, Peter Cheng, Hideo Tsukada, Edward G Robins, Julian L Goggi, Anna M. Barron

**Affiliations:** Neurobiology of Aging and Disease Laboratory, Lee Kong Chian School of Medicine, Nanyang Technological University Singapore, Singapore; Institute of Bioengineering and Bioimaging, Agency for Science, Technology and Research (A* STAR), 11 Biopolis Way, #03/07-10 Helios, Singapore, 138667; Clinical Imaging Research Centre, Yong Loo Lin School of Medicine, National University of Singapore, Singapore, 117599; Central Research Laboratory, Hamamatsu Photonics K.K., Shizuoka, Japan

**Author notes:** **Correspondence to**: Anna M. Barron, Lee Kong Chian School of Medicine, Clinical Sciences Building, 11 Mandalay Rd, Singapore 308232. **Declarations of interest:** Hideo Tsukada is employee of Hamamatsu Photonics K.K., and reagents for synthesis of ^18^F-BCPP-EF were supported in part by the company’s budget. The patents of [18F]BCPP-EF (PCT/JP2013/072442, US2015/225368A1, EP13 831 121.2, ZL201380044392.5) have been registered for Hamamatsu Photonics K.K. All other authors declare nothing to disclose.

**Keywords:** Mitochondria, Alzheimer’s disease, Positron emission tomography, Functional neuroimaging, Tau

## Abstract

Dysfunction of the energy producing organelle of the cell, mitochondria, plays a pivotal role in Alzheimer’s disease (AD). We have recently used a novel positron emission tomographic (PET) imaging tracer targeting mitochondrial complex I (MC-I) to visualize mitochondrial abnormalities in the brains of living tau transgenic (TauTg) mice. MC-I mediates the first and limiting step in oxidative phosphorylation, the primary source of neuronal energy production. Here we used MC-I-PET to test if inhibition of mutant tau expression through transgene suppression with doxycycline could reverse mitochondrial defects in a mouse model of tauopathy and evaluate the efficacy of a MC-I-targeted candidate therapeutic, Mdivi-1. We found that late-stage suppression of mutant tau did not rescue mitochondrial deficits measured *in vivo* by MC-I-PET, despite reduced burden of tauopathy and neuroinflammation. These findings demonstrate that mitochondrial dysfunction may continue even if tauopathy is halted, particularly if initiated at late-stage disease. Further, we demonstrate the potential application of MC-I-PET for monitoring therapeutic efficacy, surprisingly finding detrimental effects of the mitochondrial-targeted candidate therapeutic, Mdivi-1, in TauTg mice. These findings directly contrast with the beneficial effects of Mdivi-1 observed in other models of neurodegeneration. Together, our findings highlight the need for clinical endpoints measuring mitochondrial damage in addition to markers of tauopathy in the assessment of disease prognosis and efficacy of candidate therapeutics and demonstrates the potential application of MC-I-PET to meet this need.

## Introduction

The brain is the most energetically demanding organ of the body and is highly vulnerable to metabolic stress. Dysfunction of the energy producing organelle of the cell, mitochondria, plays a pivotal role in brain aging and neurodegenerative disease, including Alzheimer’s disease (AD). Bioenergetic and mitochondrial dysfunction are observed in early AD, worsening with disease progression (1–5). Recently, mitochondrial abnormalities have been visualised in the living brain by positron emission tomography (PET) targeting mitochondrial complex I (MC-I)

(6). MC-I carries out the first step of oxidative phosphorylation (7), which produces most neuronal energy, and is the main site of mitochondrial reactive oxygen species (ROS) production. In early AD, mitochondrial dysfunction measured by MC-I-PET was closely associated with tauopathy burden (5), a pathological hallmark of AD strongly linked to disease stage and severity. Growing evidence suggests that tauopathy, caused by intraneuronal aggregation of hyperphosphorylated forms of tau, disrupts mitochondrial function contributing to tau-mediated neurodegeneration in AD. Here we explore the use of MC-I-PET for addressing the role of mitochondrial dysfunction in tauopathy and evaluating the efficacy of mitochondrial targeted therapeutics.

In AD, decreased expression and activity of mitochondrial respiratory chain enzyme complexes are observed in vulnerable regions of the brain (8), and specific impairments in MC-I have been reported in both patients and animal models of the disease (8, 9). *In vitro* studies have shown that overexpression of human mutant tau inhibits MC-I activity and decreases ATP levels leading to reduced neuronal viability (10). Further, N-terminal fragments of tau, which are enriched in mitochondria from AD brains (11), have been shown to impair mitochondrial oxidative phosphorylation (12). Mitochondrial dysfunction is also reported in tau transgenic mouse models (TauTg), including reduced MC-I activity, impaired respiration and ATP synthesis (13), and mitochondrial mislocalization (14, 15). Similar abnormalities in neuronal mitochondrial localization are observed in postmortem AD brains (14), which is hypothesized to lead to synaptic failure and dying back of neurons (14). In a TauTg mouse expressing a human tau mutation associated with frontotemporal dementia, we observed reduced MC-I signals *in vivo* assessed using MC-I-PET in the cortex and hippocampus, colocalizing with regions of tauopathy and substantial neuronal loss (16). Interestingly, suppression of mutant tau expression in this TauTg mouse model has been shown to reverse abnormalities in mitochondrial tracking and distribution, halt neuronal loss, and result in functional recovery, despite continued development of neurofibrillary tangles (17, 18). Together, these findings suggest soluble forms of tauopathy disrupt mitochondrial function, which may play a key role in AD pathogenesis and tau-induced neurodegeneration.

Conversely, MC-I inhibition has been shown to induce tau hyperphosphorylation and neuronal death both *in vitro* and *in vivo* (19). In both cultured neurons and TauTg mice, a plant-derived MC-I inhibitor that has been implicated in the development of an environmental form of tauopathy was found to exacerbate tauopathy and induced mitochondrial mislocalization leading to neuronal death (20, 21). Likewise, the ROS-inducing, irreversible MC-I inhibitor, rotenone, also increases hyperphosphorylated tau accumulation in rat brain (22). Mechanistically, MC-I inhibition may promote tau hyperphosphorylation through ROS-mediated activation of glycogen synthase kinase-3beta (GSK3β) (23, 24). However, in contrast, some studies suggest MC-I inhibition may have a neuroprotective in AD (25–27). This apparent discrepancy may be explained by different effects of MC-I inhibitors on ROS production. MC-I inhibitors are typically classified into Class A or B inhibitors according to ROS generating properties, with class A MC-I inhibitors strongly inducing ROS production, and Class B MC-I inhibitors attenuating ROS production (28). Interestingly, a recent study employing a Class B MC-I inhibitor demonstrated improved outcomes in a mouse model of AD, including reduced amyloid accumulation and tau phosphorylation, and restoration of mitochondrial trafficking and biogenesis (26). MC-I-PET may provide a valuable translational tool for evaluation of candidate mitochondrial-targeted therapeutics *in vivo*.

In this study we use MC-I-PET to test if inhibition of mutant tau expression through suppression of transgene expression with doxycycline (DOX) could reverse mitochondrial defects in a mouse model of tauopathy and evaluate the efficacy of a MC-I-targeted candidate therapeutic, Mdivi-1. Mdivi-1 has been shown to have beneficial effects in several models of neurodegenerative disease (29–31), and was recently identified as a class B reversible MC-I inhibitor that attenuates ROS production (28, 32, 33). To understand how *in vivo* measures of mitochondrial function determined by MC-I-PET relate to underlying tauopathy, inflammation, and neuronal loss, we use *ex vivo* analyses of neuropathology in tissues from scanned mice. These studies aim to provide insight into the feasibility of the application of MC-I-PET for therapeutic monitoring and sensitivity of *in vivo* MC-I-PET signals to changes in AD pathogenesis.

## Methods

### Cell culture and transfection

Mouse neuroblastoma cell line Neuro2A (N2A) was grown and maintained in complete Dulbecco’s modified Eagle serum (DMEM) (Nacalai Tesque. #08458-45) containing 10% fetal bovine serum (FBS) (Gibco, #10270106) and 1% penicillin-streptomycin (Gibco, #15140122) (DMEM-COM) at 37 °C in 5% CO2. Cells were transfected with either pRK5-EGFP-Tau (#46904) or pRK5-EGFP-Tau P301L (#46908). Both plasmids were gifts from gift from Karen Ashe (34).(34). For transfection, media was replaced with Opti-MEM (Thermo Fisher, #31985062) 20 min prior to adding transfection mix containing 100 ng of DNA with Lipofectamine 2000 (1: 5 ratio) in Opti-MEM dropwise to cells. Transfection media was replaced with cell maintenance media (DMEM-COM) after an hour and cells were assayed within 24 hours of media replacement.

### Mitochondrial bioenergetics

Transfected N2A were plated into XFe96 cell culture plates at a density of 20,000/well in 200 μL of DMEM high glucose phenol free media (Thermo Fisher, #31053028) with 20% FBS and allowed to adhere overnight in 37°C incubator with 5% CO_2_. XF sensor cartridges were hydrated with sterile water and kept at 37°C overnight in non-CO_2_ incubator. Prior to experiments, the sensor cartridges were hydrated with 200 μL XF Calibrant and kept at 37°C in non-CO_2_ incubator for 45 – 60 min and the injection ports were then loaded with test compounds. For Mito Stress experiments, the media was removed and replaced with assay media XF RPMI medium (pH 7.4) (#103576-100) supplemented with 1 mM pyruvate (#103578-100), 2 mM glutamine (#103579-100), and 10 mM glucose (#103577-100). The cells were incubated at 37°C in non-CO_2_ incubator for 60 min prior to assay. Oxygen consumption rate (OCR) was measured during sequential injection of oligomycin (1.5 μM final concentration), Carbonyl cyanide-4 (trifluoromethoxy) phenylhydrazone (FCCP, 2 μM final concentration) and Rotenone/Antimycin A (0.5 μM final concentration).

### ROS assay

Cellular ROS of tau transfected N2A was measured using DCFDA / H2DCFDA - Cellular ROS Assay Kit (Abcam, ab113851) as per manufacturer’s protocol. Briefly, N2A were seeded in a dark, clear bottom 96-well microplate (25, 000 cells per well) and transfected with either normal or mutant human tau. Mutant human tau transfected N2A were treated with vehicle, Mdivi-1 1µM or 10µM for 5 hrs prior to assay. Cells were then incubated with DCFDA for 45 min at 37°C in the dark and end point fluorescence was measured immediately on a fluorescence plate reader at Ex/Em = 485/535 nm.

### Immunocytochemistry

N2A were seeded on glass coverslips and transfected with mutant human tau. Transfected N2A were incubated in ice-cold 4% paraformaldehyde for 10 min and washed twice in PBS. The cells were permeabilized with 0.1% Triton X-100 and blocked in 5% Bovine Serum Albumin (BSA) + 0.2% Triton X-100 for an hour. Cells were then incubated in primary antibody anti-Phospho Tau (S199/202) (Invitrogen, 44-768G) overnight at 4 °C, washed three times with PBS for 5 min each and incubated with secondary antibody Alexa Fluor® 647 (Abcam, ab150067) for an hour at room temperature. Cells were washed with PBS and mounted onto glass slides with Fluoroshield with DAPI (Abcam, ab104139).

### Animals and treatments

C57BL/6 (wild type, WT) and rTg4510 (TauTg) mice expressing a human tau mutation (P301L) on a regulatable promoter system enabling suppression of mutant tau with DOX administration (TauTg) (17) were bred and maintained under specific pathogen-free conditions at Lee Kong Chian School of Medicine Animal Research Facilities with food and water available ad libitum. All experiments were carried out in accordance with the National Advisory Committee for Laboratory Animal Research guidelines (Singapore) and approved by the by the NTU Institutional Animal Care and Use Committee (IACUC # A18041). Mice were genotyped and genotype was confirmed by presence of human tau by western blot and immunohistochemistry. For DOX study, TauTg-DOX mice were fed with 200mg/kg doxycycline hyclate (TD.00502) from 6.5 – 8 months of age as previously described (n=3-6/group) (17). For MC-I study, mini-osmotic pump (ALZET® Mini-Osmotic Pump Model 2006, DURECT Corporation, Cupertino, CA) was implanted subcutaneously in 5 month old mice on the lateral flank (n=3-4/group) to deliver vehicle (PBS) and MC-I inhibitor, Mdivi-1 (0.1 mg), over 1 month via a cannula inserted into the right lateral ventricular space. One vehicle treated mouse died during the experiment. Brains were collected from scanned mice, half fixed for immunohistochemistry (IHC), and half frozen for western blot analysis.

### In vivo PET-imaging

^18^F-BCPPEF was synthesized according to previously described methods (35, 36). Anesthetized mice (1% isoflurane via inhalation) were injected with ^18^F-BCPPEF (∼ 10MBq in 0.2 ml) via the lateral tail vein and static images were acquired at 30-50 min post injection. Body temperature and respiration rate were monitored during imaging using the Biovet physiological monitoring system. Post-analysis of reconstructed calibrated images was performed with FIJI and Amide software (version 10.3 Sourceforge). The PET images were co-registered to an MRI brain atlas to confirm the anatomical location of the radiopharmaceutical uptake. Uptake of radioactivity calculated as % of the injected dose per gram (%ID/g) of tissue was determined by the placement of volumes of interest (VOI) over the hippocampus, cortex or cerebellum. Using the cerebellum, which is devoid of neuropathology, as a reference region, the target-to-reference ratio was calculated as a percentage as previously described (37).

### Immunohistochemistry (IHC)

Fresh frozen hemibrains were sectioned in the sagittal plane at 40 μm by a cryostat. Sections were fixed in 4% paraformaldehyde and labelled using antibodies directed against ionized calcium-binding adaptor molecule-1 (IBA-1, rabbit polyclonal, WAKO 019-19741), GFAP (Invitrogen 13-0300), NeuN (Abcam ab177487), AT8 (Invitrogen MN1020). All staining, imaging, and analysis parameters were maintained consistently between samples for each marker. Images were captured using confocal microscope (Zeiss Laser Scanning Microscope-800 upright microscope) with Zeiss ZEN imaging software and analyses were performed in an experimenter-blinded manner using ImageJ. Images were background subtracted using rolling ball radius of 50 pixels prior to analysis or use in figures. Images of CA1 and cortical region were captured at ×40 magnification. NeuN-positive cells in cortical region were counted using the cell-count tool. For measurement of CA1 width, images of NeuN staining in the CA1 region, three lines per image were drawn across the CA1 region at equidistant intervals using the straight-line tool, and the average length was measured, as previously described (38). NeuN staining and CA1 width analysis was repeated three times on non-consecutive sections for each animal and an average used for analysis of group-wise differences. The sagittal brain sections were imaged using Carl Zeiss Axio Scan.Z1 slide scanner and the total area of hippocampal region was measured using Zeiss ZEN imaging software. For quantification of IBA-1, GFAP and AT8, an image of the frontal cortex and hippocampus was captured in three sections per animal and after background subtraction, 8-bit images were thresholded and percent area measured. Brightness is linearly adjusted in images presented in figures, with adjustment applied equally to all samples.

### Western blot

Hemibrains were homogenised in Tris-buffered saline (TBS) (Thermo Fisher, #28358) containing 1% Halt™ protease and phosphatase inhibitor cocktail (Thermo Fisher, 78442). Homogenates were centrifuged at 27,000×g for 20 min at 4 °C, and the supernatant was diluted in sodium dodecyl sulfate-sample buffer containing β-mercaptoethanol (2.5%) and heat-treated at 55 °C for 15 min. Samples were separated by gel electrophoresis in 12% Mini-PROTEAN® TGX™ Precast Protein Gels (Bio-Rad, 4561043) and transferred to membranes. Membranes were incubated with EveryBlot Blocking Buffer (Bio-Rad, 12010020) for 60 min at RT and were probed with anti-vinculin (Abcam, ab219649), anti-ATPB (Abcam, ab14730), anti-VDAC1/Porin (Abcam, ab186321), anti-complex I MT-ND1 (Invitrogen, 438800), Total OXPHOS Rodent WB Antibody Cocktail (Abcam, ab110413), anti-Phospho-Tau (Ser202, Thr205) Monoclonal Antibody (AT8) (Invitrogen, MN1020), anti-Tau (Tau12) (Sigma Aldrich, MAB2241), anti-Phospho Tau (S199/202) (Invitrogen, 44-768G), and anti-PSD95 (Cell Signal, 3409S). Chemiluminescent signals were detected using Radiance ECL (Azure Biosystems, AC2204) and immunoreactivity was visualized by Bio-Rad ChemiDoc™ MP Imaging System. Membranes were stripped with WB Stripping Solution Strong (Nacalai Tesque, 05677-65) for 30-45 min at 45°C, checked with Ponceau S staining solution (Alfa Aesar, J63139) and re-blocked with blocking buffer prior to re-probing. Band intensities were quantified using the NIH ImageJ software and normalized against vinculin levels. All samples were probed in at least two blots and the average of the two technical replicates was used for statistical analysis.

## Results

### Suppression of mutant tau does not rescue mitochondrial deficits in TauTg mice measured by MC-I-PET

Expression of frontotemporal dementia associated P301L mutant human tau (mTau) caused defects in mitochondrial localization and function in cultured N2a neuroblastoma cells (Fig. 1). mTau expression was associated with mitochondrial clustering around the nucleus (Fig. 1C) and impaired mitochondrial respiration measured by oxygen consumption rate (OCR) in the Seahorse Mitostress test (Fig. 1D, E). This included reduced basal respiration, maximal respiration and ATP production in mTau expressing N2a cells (*t*-test: basal respiration: *t* = 3.44, *p* = 0.0064; maximal respiration: *t* = 4.65, *p =* 0.0009; ATP production: *t* = 3.35, *p* = 0.0073; Fig.1D-E). Since we have previously shown that TauTg mice expressing this mutant form of human tau exhibit mitochondrial dysfunction from early stages of neuropathology (16), we hypothesized that suppression of mutant tau could reverse mitochondrial deficits detected *in vivo* using MC-I-PET. To test this, tauopathy was suppressed from 6 months of age for a period of six weeks via treatment with DOX (Fig. 2A). This was designed to test if an intervention that reduces toxic forms of soluble tau could attenuate mitochondrial defects if initiated after formation of neurofibrillary tangles and neuronal loss, modelling symptomatic late-stage disease.

**Fig. 1.**
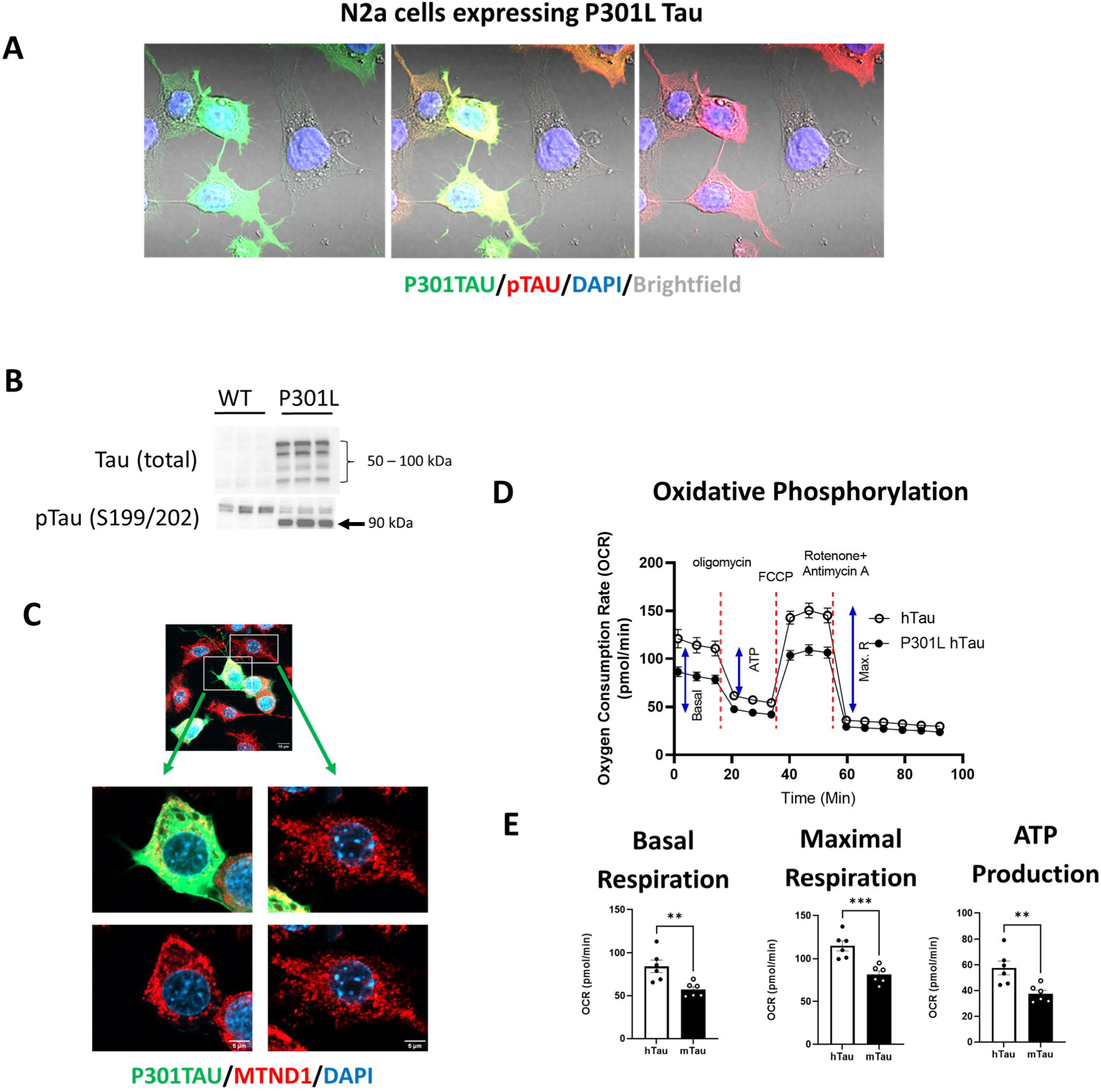
Mutant P301L tau induces mitochondrial dysfunction *in vitro*. **(A)** Mouse N2a neuroblastoma cells expressing mutant P301L human tau (mTau, green). Phosphorylated tau (ser199/202; red) immunoreactivity observed in mTau expressing cells. Cells counterstained with DAPI (blue) to show nucleus and overlaid on brightfield image (grey). (**B**) Representative western blot showing total tau and phosphorylated tau in N2a cells expressing mTau. (**C**) Representative image of mouse N2a neuroblastoma cells expressing mTau (green) with mitochondrial complex I immunoreactivity (MTND1, red). We observed mitochondrial clustering around nucleus of cells expressing mutant hTau (left) compared to cells not expressing mutant hTau (right) with mitochondria more evenly distributed across cell body. (**D**) Seahorse Mitostress test of N2a cells expressing normal human tau (hTau) versus mTau showing mean ± SEM oxygen consumption rate (OCR). Basal respiration (Basal), ATP production (ATP) and maximal respiration (Max. R) indicated on WT curve. ATP synthase inhibitor, oligomycin was injected to inhibit ATP-linked mitochondrial respiration as a measure of mitochondrial ATP production. The mitochondrial uncoupler, FCCP was injected to determine the maximal respiration rate. To completely inhibit mitochondrial respiration and determine contribution of non-mitochondrial respiration to the measured OCR, mitochondrial complex I and III was inhibited by injection of a mixture of rotenone and antimycin. (n = 6 wells/group). (**E**) Quantification of basal respiration (*left)*, maximal respiration (*middle*) and ATP production (*right)* calculated from Mitostress assay. \*\**p* < 0.01, \*\*\**p* < 0.001.

**Fig. 2.**
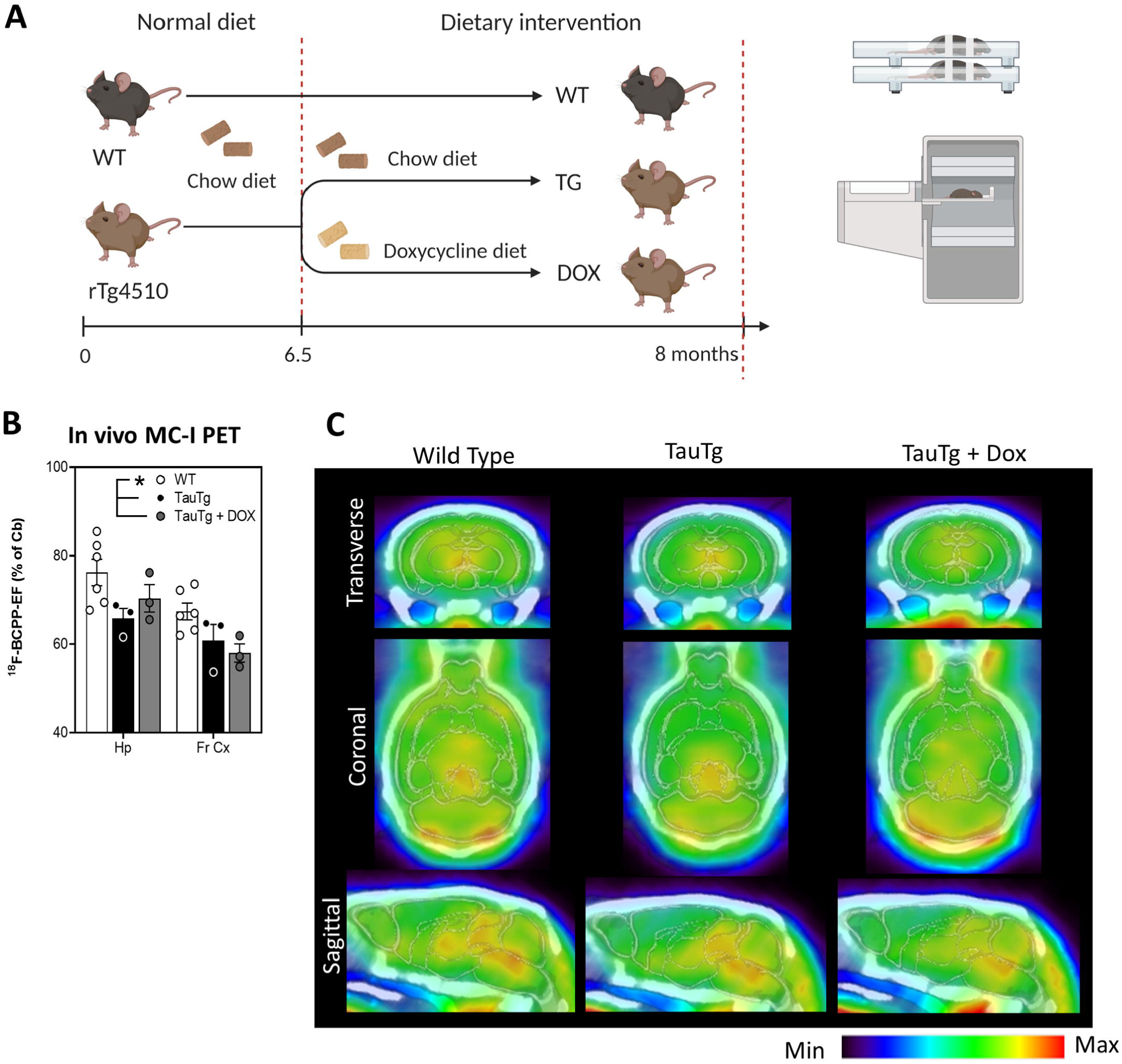
Late-stage, short-term suppression of tauopathy does not rescue MC-I deficits detected by ^18^F-BCPP-EF-PET in TauTg mice. (**A**) Schematic of experimental design. Mutant tau was supressed in TauTg mice for a period of 6 weeks until 8 months of age via DOX feeding, then mitochondrial function assessed *in vivo* by MC-I-PET. At completion of experiment the brains were collected from scanned mice for biochemical analysis of tauopathy, mitochondrial markers, neuronal loss and neuroinflammation. (**B**) Average ratio of ^18^F-BCPP-EF uptake (relative cerebellum, cb) demonstrating reduced hippocampal (Hp) and cortical (Cx) MC-I signals in TauTg mice even after DOX treatment. (**C**) Representative coronal and sagittal images of ^18^F-BCPP-EF distribution in 8-month-old WT and TauTg mice +/-DOX treatment. ^18^F-BCPP-EF uptake overlaid on individual MR data. \**p* < 0.01.

MC-I-PET signals were significantly reduced in the hippocampus and cortical regions of TauTg mice, but inhibition of mutant tau expression did not result in any measurable improvement in mitochondrial function in the hippocampal or cortex assessed by MC-I-PET (two-way ANOVA, main effect: treatment F_(2,18)_ = 26.71, *p* = 0.0091; FDR corrected pairwise comparisons, WT vs TauTg or TauTg + DOX: *p =* 0.0074; TauTg vs TauTg + DOX: *p* = 0.282, Fig. 2B, C). Reduced levels of TBS-soluble phosphorylated forms of 50-60 kDa tau (Ser199, Ser202) were confirmed in DOX treated animals by western blot (one way ANOVA: F = 18.1, *p* = 0.0007, FDR corrected pairwise comparisons, WT vs TauTg: *p* = 0.0002; TauTg vs TauTg + DOX: *p =* 0.0024; Fig. 3A, B). Although soluble forms of phosphorylated tau were reduced, neurofibrillary tangle formation measured by AT8 immunoreactivity was not affected by mutant tau suppression (Fig. 3C). This was consistent with previous findings in the TauTg model (17). Likewise, despite reduced levels of phosphorylated forms of tau, suppression of mutant tau did not rescue brain mitochondrial density measured by western blot using the mitochondrial marker ATPB (Fig. 3B, D), or hippocampal neuronal density measured by NeuN immunohistochemistry (one way ANOVA: ATPB: F = 5.51, *p* = 0.027, FDR corrected pairwise comparisons, TauTg + DOX vs WT *p =* 0.02, TauTg + DOX vs TauTg *p* = 0.12; PSD-95: F = 8.4, *p* = 0.009; NeuN: F = 8.34, *p =* 0.009, FDR corrected pairwise comparisons WT vs TauTg or TauTg + DOX: *p* < 0.022; TauTg vs TauTg + DOX: *p =* 0.99; Fig. 3E, F). *In vivo* MC-I PET signals negatively correlated neurofibrillary tangle burden (r = - 0.713, *p* = 0.012) and positively correlated with hippocampal NeuN neuronal density (r = 0.678, *p* = 0.018).

**Fig. 3.**
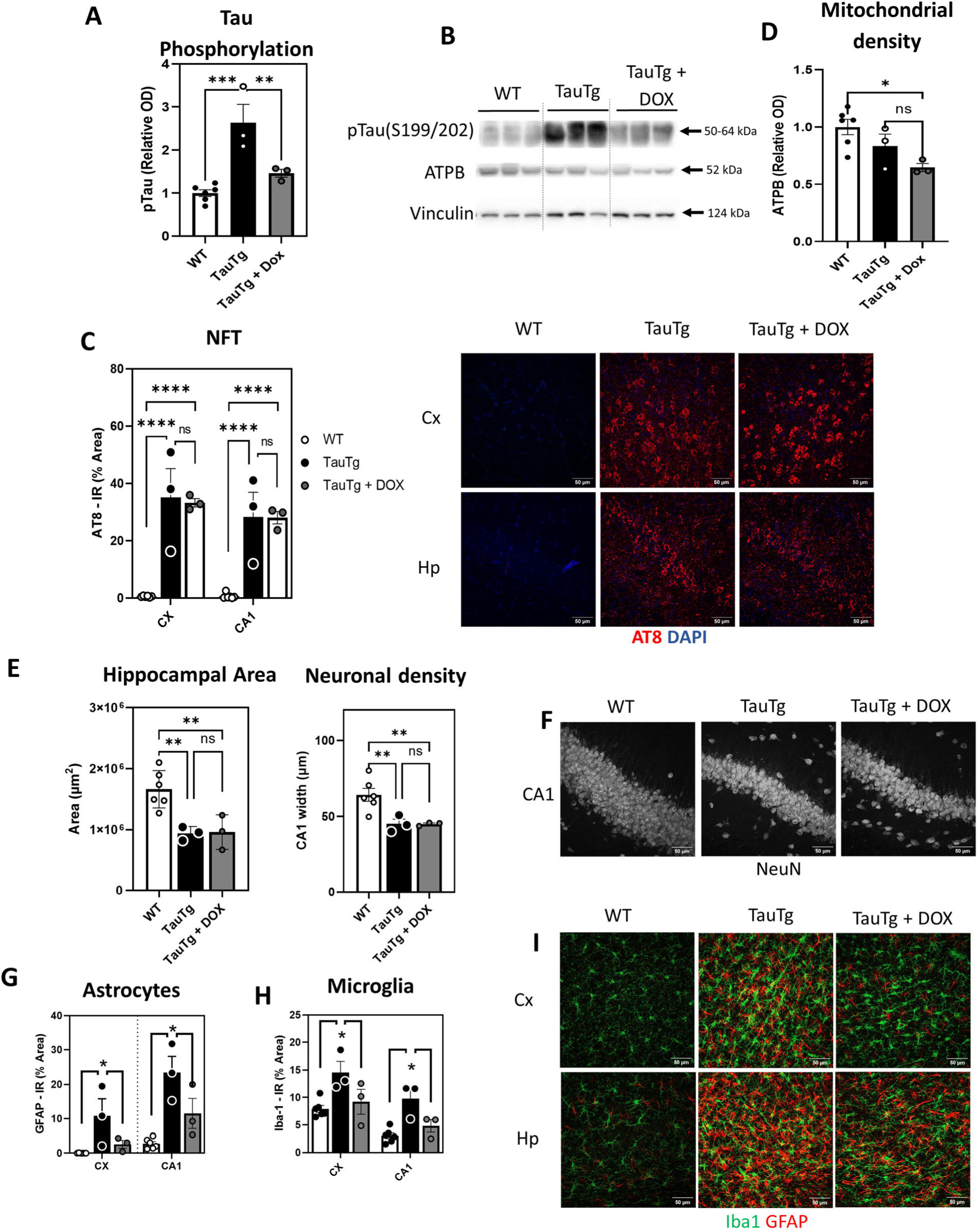
Late-stage, short-term suppression of tauopathy reduces inflammation but neurodegeneration in TauTg mice. (**A**) Quantification of 50-60kDa phosphorylated tau (Ser199, Ser202) detected by western blot. (**B**) Representative immunoblot of brain homogenate probed for phosphorylated tau (pTau), ATPB, and loading control, vinculin. (**C**) Quantification of neurofibrillary tangle marker, AT8 immunoreactivity in hippocampus (Hp) and cortical (Cx) regions (*left)* and representative confocal images of AT8 immunoreactivity in hippocampal (Hp) and cortical (Cx) regions (*right*). (**D**) Quantification of mitochondrial marker, ATPB, detected by western blot. (**E**) Quantification of hippocampal area (*left)* and hippocampal CA1 neuronal density (*right)* measured by immunohistochemistry using neuronal marker, NeuN. (**F**) Representative confocal images of NeuN immunoreactivity in hippocampal CA1 region (Hp) of WT and TauTg mice ± DOX treatment. (**G-I**) Quantification of astrocyte marker GFAP (**G**) and microglial marker IBA-1 (**H**) immunoreactivity in hippocampal and cortical regions. (**I**) Representative confocal images of IBA-1 (green) and GFAP (red) immunoreactivity. Mean ± SEM. n = 3-6/group. \**p* < 0.05, \*\**p* < 0.01, \*\*\**p* < 0.001, \*\*\*\**p* < 0.0001.

Whilst suppression of tauopathy failed to rescue mitochondrial and neuronal loss, it did reduce markers of neuroinflammation. Reduced immunoreactivity of the astrocyte marker GFAP and microglial marker IBA-1 were detected in hippocampal and cortical regions of DOX treated TauTg mice (two way ANOVA, main effect: treatment: GFAP F _(2, 18)_ = 55.27, *p* < 0.0001, FDR corrected pairwise comparisons WT vs TauTg *p* < 0.0001, TauTg vs TauTg + DOX *p* = 0.0031; IBA-1 F_(2,18)_ = 43.07, *p* = 0.0001, FDR corrected pairwise comparisons WT vs TauTg *p* < 0.0001, TauTg vs TauTg + DOX *p* = 0.002; Fig 3G - I). Markers of neuroinflammation were strongly correlated with levels of phosphorylated tau (GFAP: r = 0.916, *p* > 0.0001; IBA-1: r = r = 0.657, *p* = 0.02).

### *In vitro* characterization of MC-I inhibitor, Mdivi-1, in tauopathy model

Here we examined the effect of a mitochondrial-targeted candidate therapeutic, Mdivi-1 on cellular bioenergetics and mitochondrial morphology in N2a neuroblastoma cells expressing P301L mutant tau (mTau). Initially identified as a candidate inhibitor of mitochondrial fission, a recent study has demonstrated that Mdivi-1 reversibly inhibits MC-I activity in neurons(33). We confirmed this action, demonstrating Mdivi-1 dose-dependently inhibits basal respiration, maximal respiration and ATP production in neuroblastoma cells expressing mTau (Kruskal-Wallis test, basal respiration: *H* = 18.98, *p* = 0.0003, FDR corrected post-hoc compared to vehicle 1µM *p =* 0.0358, 10µM *p* = 0.0006; maximal respiration: *H* = 17.93, *p* = 0.0005, FDR corrected post-hoc compared to vehicle 1µM *p =* 0.0262, 10µM *p* = 0.0026; ATP production: *H* = 18.98, *p* = 0.0003, FDR corrected post-hoc compared to vehicle 1µM *p =* 0.0358, 10µM *p* = 0.0006; Fig. 4A, B). However, unlike the toxic and irreversible MC-I inhibitor, rotenone, Mdivi-1 reduced reactive oxygen species production, reversing mTau associated increases in ROS production in the N2a cells (ROS, Kruskal-Wallis test: *H* = 11.73, *p* = 0.0084, FDR corrected post-hoc, hTAU vs mTAU *p =* 0.016, mTAU vehicle vs 1µM *p =* 0.3, vs 10µM *p* = 0.016; Fig. 4C). Surprisingly, Mdivi-1 did not induce any noticeable changes in mitochondrial morphology in N2a cells expressing mTau, (Fig. 4D). No significant effect of Mdivi-1 treatment was observed on mitochondrial number (*t-*test, *t* = 1.06, *p* = 0.29, n = 12-22 individual cells) or footprint (*t-*test, *t* = 0.82, *p* = 0.42, n = 12-22 individual cells). This was consistent with other recent findings indicating Mdivi-1’s inhibition of respiration may not be directly related to the inhibitory effects of Mdivi-1 on Drp1 and mitochondrial fission (33). We also found that treatment of cells with Mdivi-1 did not induce cell death (<1% dead cells in all conditions – vehicle, 1 & 10µM Mdivi-1).

**Fig. 4.**
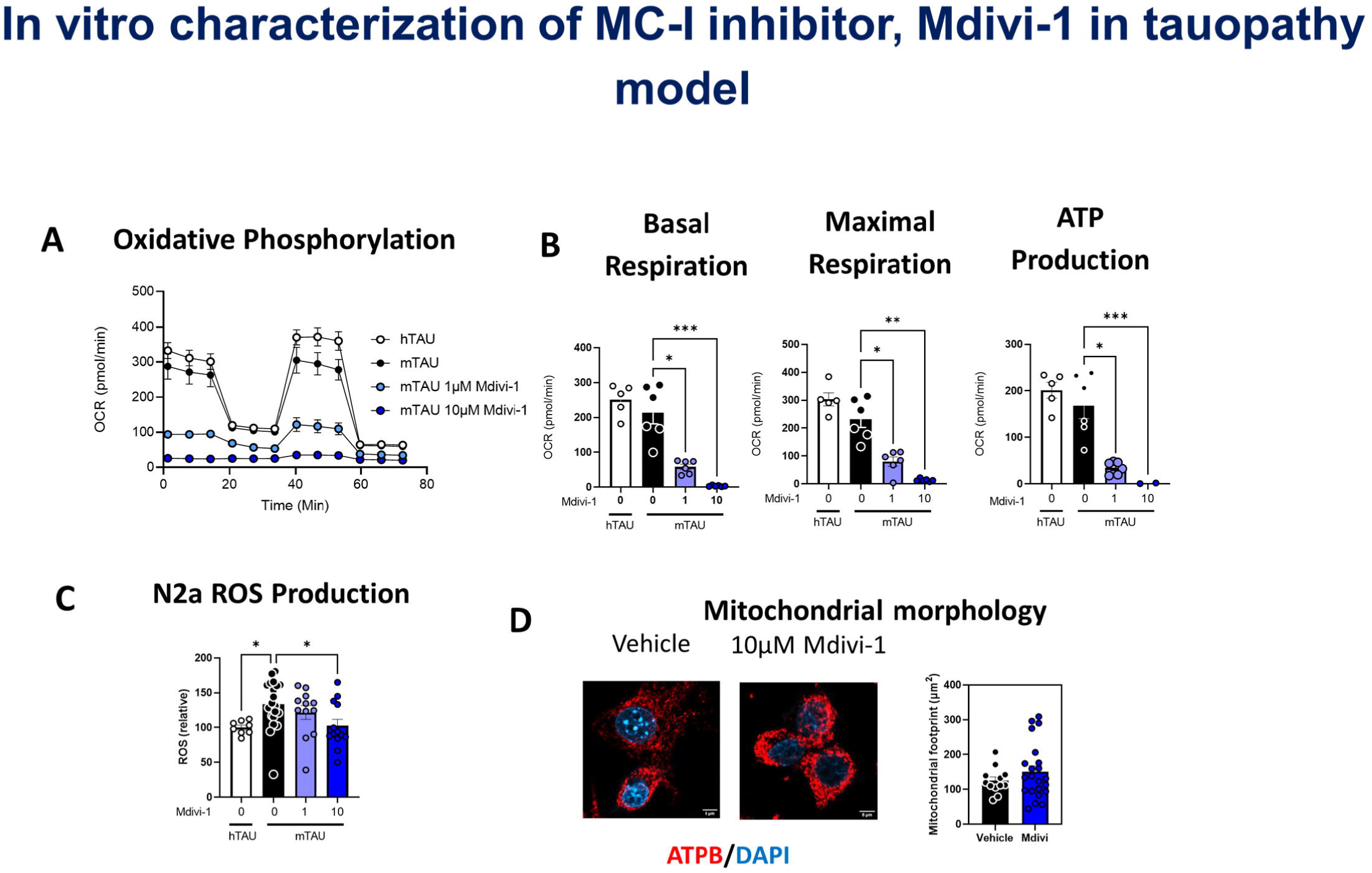
MC-I inhibitor, Mdivi-1, reduces mitochondrial respiration while reversing mutant tau-induced ROS production *in vitro*. **(A)** Seahorse Mitostress test of N2A transfected with P301L mutant human Tau (mTau) and treated with vehicle 0.1µM and 10µM Mdivi-1. (**B**) Quantification of basal respiration, maximal respiration and ATP production calculated from Mitostress assay showing Mdivi-1 dose-dependently inhibited N2a bioenergetics. (**C**) Expression of mTau was associated with increased ROS production in N2a cells. Mdivi-1 dose-dependently reduced ROS levels in N2a cells expressing mTau. (**D**) Representative images mitochondrial marker, ATPB (red) immunoreactivity counterstained with DAPI (blue) in N2a cells expressing mTau treated with either vehicle or 10µM Mdivi-1. Quantification of mitochondrial footprint (*right*, n = 12 -22 cells/treatment). \**p* < 0.05, \*\**p* < 0.01, \*\*\**p* < 0.001.

### Mdivi-1 inhibits *in vivo* MC-I-PET signals and worsens pathology in TauTg mice

Next we investigated the effect of Mdivi-1 treatment on *in vivo* mitochondrial deficits in TauTg mice measured by MC-I-PET (Fig. 5A). Based on our *in vitro* characterization of Mdivi-1, mice were treated with 5nmol/day, calculated to yield 10µM Mdivi-1 in CSF, which would result in an order of magnitude lower concentrations in the brain parenchyma (based on 40µL total CSF volume, 350nL/min turnover (39). MC-I-PET signals were signficantly reduced in hippocampal and cortical regions of mice intracerebroventricularly infused with Mdivi-1 over 1 month (two-way ANOVA, treatment main effect, F_(1, 15)_18.32, *p* = 0.009; Fig. 5B, C). This corresponded to reduced mitochondrial density in the brains of Mdivi-1 treated mice measured by western blot using the mitochondrial marker ATPB (*t*-test: *t =* 2.58, *p* = 0.049; Fig. 5D, E). Mdivi-1 treatment also resulted in an increase in 64 kDa phosphorylated tau species detected by AT8 (Ser202/Thr 205; *t*-test: *t* = 2.63, *p =* 0.046), but not the smaller 50 – 60kDa ser199/202 phosphorylated tau species (t-test: *t = 0.64, p* = 0.55; Fig. 5D, F, G). This longer, 64kDa phosphorylated species of tau has been previously shown to increase with disease progression in the tauTg mouse (40). Hippocampal area was also decreased following Mdivi-1 treatment (*t-*test: *CI* = -6.3e+05, -1.3e+05*, p* = 0.031), although no significant effect of treatment was observed on hippocampal neuronal density or markers of neuroinflammation (Fig. 5H-K). These findings indicate that Mdivi-1 exacerbates neuropathology progression in TauTg mice, in contrast to therapeutic benefits observed in other models of neurodegenerative disease (29, 41, 42).

**Fig. 5.**
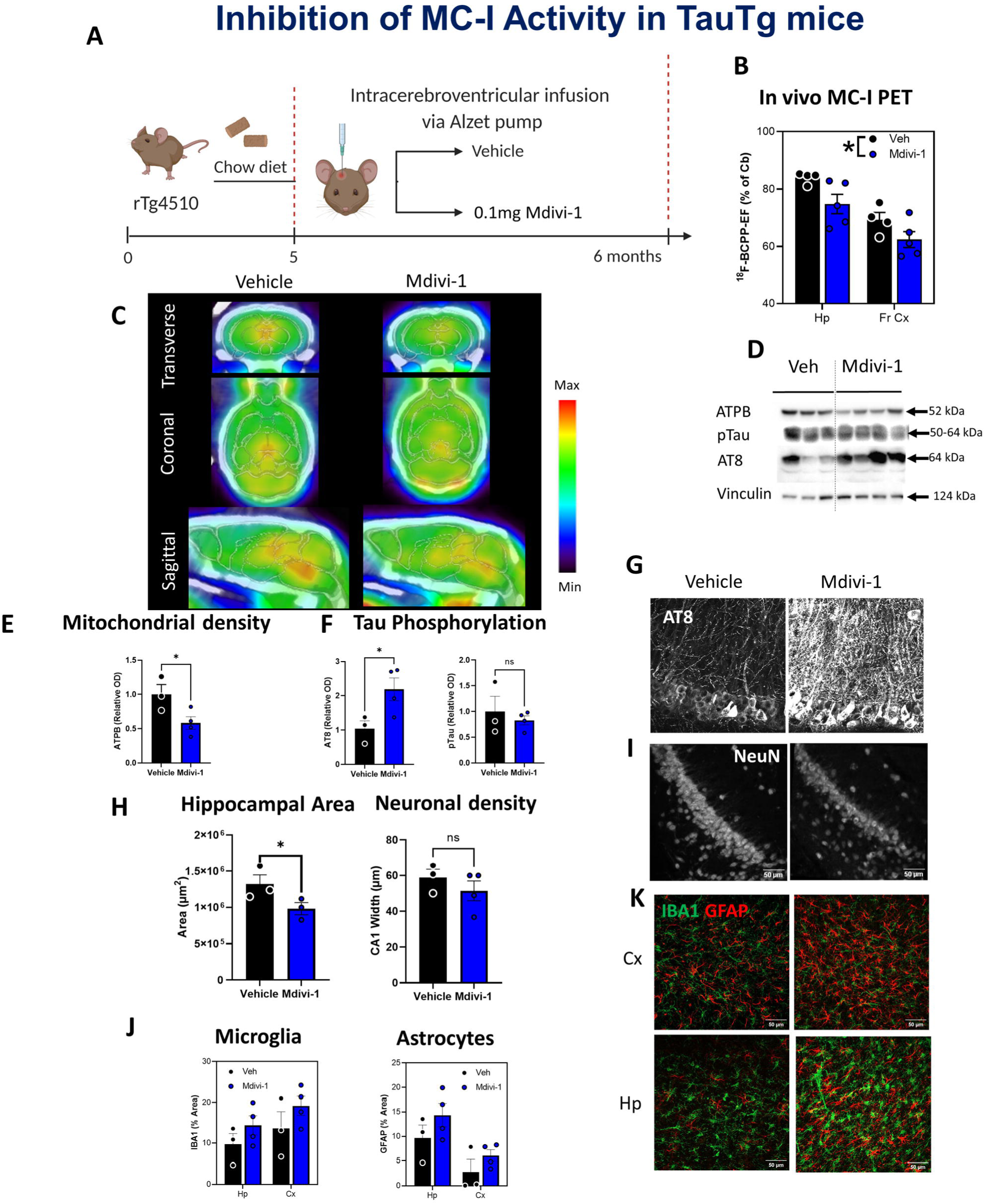
MC-I inhibitor, Mdivi-1, impairs *in vivo* MC-I-PET signals and worsens tauopathy in TauTg mice. (**A**) Schematic of experimental design. TauTg mice were treated with Mdivi-1 (0.1mg) or vehicle from 5 months of age for 1 month, then mitochondrial function was assessed *in vivo* by MC-I-PET. (**B)** Average ratio of ^18^F-BCPP-EF uptake (relative cerebellum, cb) demonstrating reduced hippocampal (Hp) and cortical (Cx) MC-I signals Mdivi-1 treated mice. (**C**) Representative coronal and sagittal images of ^18^F-BCPP-EF distribution in TauTg mice treated with vehicle or Mdivi-1 0.1mg via intracerebroventricular infusion. ^18^F-BCPP-EF uptake overlaid on individual MRI data. (**D**) Representative immunoblot of brain homogenate probed for ATPB, 50-60kDa phosphorylated tau (ser199/202), 64 kDa phosphorylated tau (AT8), and loading control, vinculin. (**E**) Quantification of mitochondrial marker, ATPB, detected by western blot. (**F**) Quantification of 64kDa (AT8, *left)* and 50-60kDa (ser199/202, *right*) phosphorylated tau measured by western blot. (**G**) Representative confocal images of AT8 immunoreactivity in hippocampus. (**H**) Quantification of hippocampal area (*left)* and neuronal density (*right)* measured by immunoreactivity of the neuronal marker, NeuN. (**I**) Representative confocal images of NeuN immunoreactivity in hippocampal CA1 region (Hp) of vehicle and Mdivi-1 treated TauTg mice. (**J-K**) Quantification of microglial marker IBA-1 (**J**) and astrocyte marker GFAP (**K**) immunoreactivity in hippocampal and cortical regions of vehicle and Mdivi-1 treated TauTg mice. (**L**) Representative confocal images of IBA-1 (green) and GFAP (red) immunoreactivity in hippocampal (Hp) and cortical (Cx) regions. Mean ± SEM. n = 3-4/group. \**p* < 0.05.

## Discussion

Here we used MC-I-PET to test if inhibition of mutant tau expression could reverse mitochondrial defects in a mouse model of tauopathy and evaluate the efficacy of a MC-I-targeted candidate therapeutic, Mdivi-1. We found that late-stage suppression of mutant tau did not rescue mitochondrial deficits measured *in vivo* by MC-I-PET, despite reduced burden of tauopathy and neuroinflammation, which was corroborated by *ex vivo* analyses. Importantly, our findings demonstrate that mitochondrial dysfunction may continue even if tauopathy is halted, particularly if initiated at late-stage disease, findings which have important potential implications for the development of tau-targeted therapies. Further, we demonstrate the potential application of MC-I-PET for monitoring therapeutic efficacy, surprisingly finding detrimental effects of the mitochondrial-targeted candidate therapeutic, Mdivi-1, in TauTg mice. These findings directly contrast with the beneficial effects of Mdivi-1 observed in other models of neurodegeneration (29–31). Together, our findings highlight the need for clinical endpoints measuring mitochondrial damage in addition to markers of tauopathy in the assessment of disease prognosis and efficacy of candidate therapeutics and demonstrates the potential application of MC-I-PET to meet this need.

In this study, we used brief (6 weeks) tau suppression at an advanced stage of pathology (6 months of age), characterized by extensive NFT formation, neuronal loss and inflammation throughout forebrain, including the hippocampus frontal cortex (17). Although suppression of mutant tau reduced phosphorylated forms of 50-60 kDa tau (Ser199, Ser202), it did not halt NFT formation. This is consistent with previous findings showing NFT formation continues until old age if mutant tau suppression is initiated beyond 4 months of age (17). Likewise, although we observed reduced levels of phosphorylated forms of tau and inflammation, this did not rescue neuronal or mitochondrial loss. Interventions implemented at a younger age have demonstrated the ability to reduce the severity of NFT pathology in the forebrain (17, 43, 44), a strategy which could potentially also mitigate mitochondrial dysfunction. These findings highlight the importance of intervening at an early stage to potentially slow down the progression of NFT pathology and mitochondrial dysfunction in this model.

Since DOX has been demonstrated to suppress microglial activation (45, 46), we cannot exclude the possibility that the DOX treatment acted directly on glia to reduce markers of inflammation. However, tauopathy was strongly correlated with markers of inflammation, supporting the interpretation that reduced inflammation was a consequence of reduced levels of toxic forms of tau. Since tau-induced inflammation is thought to play a key role in mediating neurodegeneration (47, 48), we were surprised to find neuronal loss was not mitigated by tau suppression in our experiments. However, we found that *in vivo* mitochondrial signals measured by MC-I-PET were strongly associated with measures of neuronal density, supporting the notion mitochondrial dysfunction is intricately linked with tau-mediated neurodegeneration.

We then directly tested the hypothesis that mitochondrial dysfunction drives tau-mediated neurodegeneration, assessing the efficacy of the candidate therapeutic, Mdivi-1, in tauTg mice. Previous studies have shown neuroprotective effects of Mdivi-1 in both *in vitro* and *in vivo* models of neurodegeneration. For example, Mdivi-1 ameliorated mitochondrial dysfunction, rescued synaptic loss and cognitive impairment, and decreased inflammation in models of beta amyloid induced neuropathology (41, 42). Likewise, Mdivi-1 reduced hypoxia-induced mitochondrial fission and amyloid-beta accumulation (29), and reduced α-synuclein aggregation, oxidative stress and neurodegeneration in a rat model of Parkinson’s disease (30, 41, 42). However, to our surprise, in contrast to the rescue effects of Mdivi-1 in other studies, we found Mdivi-1 to worsen outcomes in a mouse model of tauopathy. Mdivi-1 treatment in TauTg mice exacerbated NFT formation, reduced mitochondrial density and worsened hippocampal atrophy. Of note, TBS-soluble forms of phosphorylated tau were not affected by Mdivi-1 treatment. These findings may suggest a link between NFT formation and mitochondrial dysfunction in this mouse model.

Mechanistically, Mdivi-1 rapidly and reversibly binds MC-I, inhibiting the electron transport chain, impairing respiration while also inhibiting ROS production (33). This is proposed to induce a mitohormetic effect that promotes neuronal survival. This was supported by our *in vitro* studies, where Mdivi-1 treatment dose-dependently reduced mitochondrial respiration while reversing tau-associated ROS production. Mdivi-1 has also been implicated in mitochondrial fission, acting as an inhibitor of dynamin-related protein 1 (Drp1), reducing the overall number of mitochondria while promoting elongation of individual mitochondria (49). However, more recently, this has been questioned in mammalian systems (33). Supporting this, we did not observe any effect of Mdivi-1 treatment on mitochondrial number or content in cultured neuroblastoma cells.

In a recent study, it was demonstrated that dysfunction of MC-I and the consequent rise in mitochondrial ROS contribute to an elevation in tau phosphorylation and aggregation when cells experience mitochondrial stress (50). Interestingly, the researchers observed that treatment with low concentrations of ROS scavengers had the most significant impact on reducing tau dimerization, indicating the possibility of mitohormetic effects, wherein a specific level of ROS demonstrates beneficial effects (50). Recent study using CRISPR screening revealed new roles of tau proteostasis in neurons and supported the role of defective mitochondria in increasing tau oligomerisation and promoting proteasomal misprocessing of tau proteins. Knocking down genes for proteins associated with electron transport chain function was shown to increase ROS level as well as levels of tau oligomers (51). Similarly, another research group showed that mitochondrial function plays a role in tau inclusion and propagation. They found that inhibition of several components of MC-I reduces tau propagation, potentially via a reduction of neuronal activity due to decreased mitochondrial function (52). Other studies targeting MC-1 have also observed beneficial effects in models of AD (27). For example partial inhibition of MC-I by small molecule CP2 was found to be beneficial in restoring mitochondrial dynamics and enhances communication between mitochondria and the endoplasmic reticulum (ER) (53), and yielded positive outcomes in reducing levels of phosphorylated tau in the presence of amyloid-beta in an AD mouse model (54). MC-I inhibitor, C458, triggered a mitochondrial stress response, leading to an increase in mitochondrial biogenesis, antioxidant signaling, and cellular energy levels, effectively mitigating cognitive decline in an AD model (27). Possible reasons for the discrepancy between these previous studies and our current findings include differences in dose, disease severity at intervention and model of disease. Interestingly, the partial MC-I inhibitor, metformin, has also been found to elicit disparite effects on neurodegeneration. For example, metformin exacerbated neurodegeneration in a model of Parkinsons disease (55), exacerbated tauopathy in TauTg mice (56), but reduced beta amyloid deposition in transgenic AD mice (57). This highlights the need for clinical endpoints measuring mitochondrial damage in the assessment of the efficacy of candidate therapeutics and our findings demonstrate the potential application of MC-I-PET to meet this need.

^18^F-BCPP-EF is the first MC-I-PET probe with high and rapid brain uptake, minimal background signal, and sustained retention within the target region, providing suitable specificity and kinetics for imaging mitochondrial function in the brain (6, 58). By imaging MC-I activity, this unique probing technique offers a deeper understanding of energy metabolism, specifically targeting the OXPHOS pathway. BCPP-EF binds to the MC-I site shared with the class A, ROS producing inhibitor, rotenone (59). Meanwhile, the candidate therapeutic tested here, Mdivi-1, inhibits MC-I activity via a separate mechanism, thought to bind to the Q site on MC-I (33). Using ^18^F-BCPP-EF, MC-I impairment has been shown to be associated with tau deposition in animal model (16) and AD patients (60, 61). A recent study using ^8^F-BCPP-EF highlighted a change in mitochondrial activity in the early stages of an AD mouse model and a dynamic shift in mitochondria corresponding to the advancement of pathology, mirroring analogous patterns observed in the early stages of the AD spectrum disorder in humans (62). ^18^F-BCPP-EF has also shown promising results in preclinical studies and proposed to be useful for monitoring MC-I function in humans in the future for the evaluation of AD pathophysiology. A recent study using MC-I-PET provided valuable *in vivo* evidence suggesting extensive cellular stress and bioenergetic abnormalities in the early stages of AD, with ^18^F-BCPP-EF evaluated to be the most sensitive longitudinal marker of disease progression. They reported a decrease in MC-I binding in AD patients, especially in the hippocampal, caudate, and thalamic region and a loss of metabolic functional reserve as the disease progresses (63). Buiding on these studies, our findings support the potentail application of MC-I-PET for treatment monitoring. One potential limitation of our approach is that differences in regional cerebral blood flow (rCBF) can alter PET-tracer uptake and confound interpretation of *in vivo* differences in PET signals. However, the reversible-type kinetics of ^18^F-BCPP-EF reduces sensitivity to rCBF changes with studies supporting the notion that ^18^F-BCPP-EF is less rCBF dependent (64, 65), thus minimizing this type of error. Further, *ex vivo* analyses, which are not susceptible to rCBF, corroborated our *in vivo* findings, confirming that MC-I-PET signals were tightly coupled to mitochondrial density and neuronal loss.

Based on the recent perspectives on the key role of mitochondrial metabolism and bioenergetics in AD, mitochondrial-targeted neuroimaging tools for diagnostic, prognostic and therapeutic monitoring are needed. Here we demonstrate the application of ^18^F-BCPP–EF as a functional marker of MC-I function and evaluated the efficacy of Mdivi-1 in TauTg mouse model of AD. Our findings highlight the potential application of MC-I-PET for evaluation of novel mitochondrial targeted therapeutics. These findings will aid interpretation of trials testing the MC-I-PET tracer in humans, enabling understanding of what *in vivo* signals indicate about mitochondrial function and Alzheimer’s pathogenesis.

## Acknowledgements

This project was funded by the Ministry of Education, Singapore, under its Academic Research Fund Tier 1 (Project ID RG42/18; AMB), the Alzheimer’s Association (AARG-18-566427 to A.M.B) and the Nanyang Assistant Professorship Award from Nanyang Technological University Singapore (AMB). The authors thank Daniel Ch’ng Yu Sheng for his pilot analysis of inflammatory markers.

## CRediT authorship contribution statement

**Jia Hui Wong:** Methodology, Investigation, Validation, Formal Analysis, Visualization, Writing – original draft. **Anselm S. Vincent:** Methodology, Investigation, Validation. **Shivashankar Khanapur:** Methodology, Investigation. **Tang Jun Rong:** Methodology, Investigation. **Boominathan Ramasamy:** Methodology, Investigation. **Siddesh Hartimath:** Methodology, Investigation, Visualization. **Peter Cheng:** Methodology, Investigation. **Hideo Tsukada:** Resources, Methodology, Writing – review & editing. **Edward G Robins:** Methodology, Investigation, Writing – review & editing. **Julian L Goggi:** Methodology, Investigation, Formal Analysis, Writing – review & editing. **Anna M. Barron**: Conceptualization, Methodology, Investigation, Formal Analysis, Visualization, Writing - original draft, Supervision, Funding acquisition.

## Notes

### Competing Interest Statement

Hideo Tsukada is employee of Hamamatsu Photonics K.K., and reagents for synthesis of 18F-BCPP-EF were supported in part by the companys budget. The patents of [18F]BCPP-EF (PCT/JP2013/072442, US2015/225368A1, EP13 831 121.2, ZL201380044392.5) have been registered for Hamamatsu Photonics K.K..

