## Supplemental Figures for "Mitochondrial complex I as a diagnostic and therapeutic target in a mouse model of tauopathy"

5 d

1 d

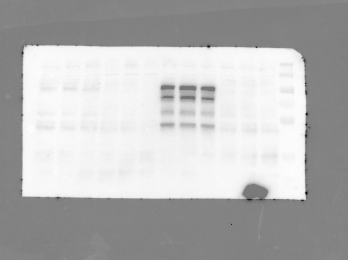

WT

P301L

WT

P301L

250 🡪

100🡪

55🡪

35🡪

Fig. S1. Raw image of representative western blot data shown in Fig 1B (top). Entire blot probed for total tau (50-100 kDa). Lane 1: 35 – 250kDa ladder. Lane 2-4 & 8-10: untransfected BV2 samples. Lane 5-7 & 11-13: mutant P301 human tau transfected BV2 samples. Lane 2-7: 1 day post-transfection, Lane 8-13: 5 days post-transfection. 1 replicate = 1 well.

5 d

1 d

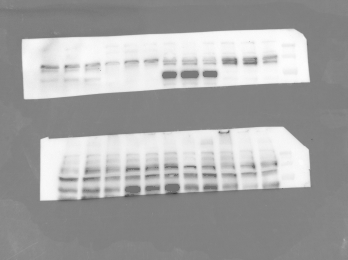

WT

P301L

250 🡪

100🡪

WT

P301L

Fig. S2. Raw image of representative western blot data shown in Fig 1B (bottom). Blot stripped, cut at approx. 60kDa and the top half reprobed for pTau (S199/202) (90 kDa). Lane 1: 35 – 250kDa ladder. Lane 2-4 & 8-10: untransfected BV2 samples. Lane 5-7 & 11-13: mutant P301 human tau transfected BV2 samples. Lane 2-7: 1 day post-transfection, Lane 8-13: 5 days post-transfection. 1 replicate = 1 well.

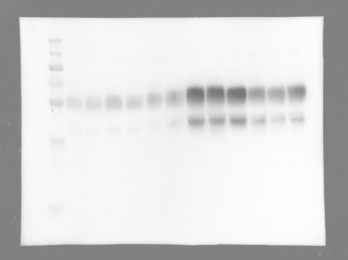

250 🡪

100🡪

55🡪

35🡪

WT

TauTg

TauTg + DOX

Fig. S3. Raw image of representative western blot data shown in Fig 3B (top). Entire blot probed for pTau (S199/202) (50-64 kDa). Lane 1: 35 – 250kDa ladder. Lane 2-7: WT mouse brain homogenate. Lane 8-10: TauTg mouse brain homogenate. Lane 11-13: TauTg + DOX treated mouse brain homogenate. Biological replicates.

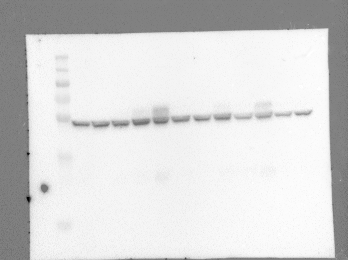

TauTg + DOX

TauTg

WT

250 🡪

100🡪

55🡪

35🡪

Fig. S4. Raw image of representative western blot data shown in Fig 3B (middle). Entire blot probed for ATPB (52 kDa). Lane 1: 35 – 250kDa ladder. Lane 2-7: WT mouse brain homogenate. Lane 8-10: TauTg mouse brain homogenate. Lane 11-13: TauTg + DOX treated mouse brain homogenate. Biological replicates.

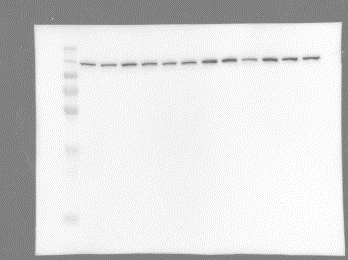

250 🡪

100🡪

55🡪

35🡪

TauTg

WT

TauTg + DOX

Fig. S5. Raw image of representative western blot data shown in Fig 3B (bottom). Entire blot stripped and reprobed for vinculin (124 kDa). Lane 1: 35 – 250kDa ladder. Lane 2-7: WT mouse brain homogenate. Lane 8-10: TauTg mouse brain homogenate. Lane 11-13: TauTg + DOX treated mouse brain homogenate. Biological replicates.

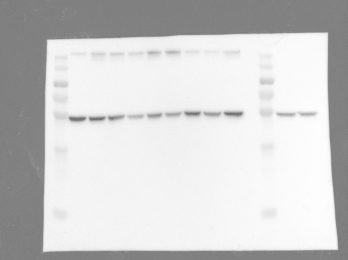

Mdivi-1

Veh

250 🡪

100🡪

55🡪

35🡪

Fig. S9. Raw image of representative western blot data shown in Fig 5D (top). Entire blot probed for ATPB (52 kDa). Lane 1: 35 – 250kDa ladder. Lane 2-4: Vehicle treated TauTg mouse brain homogenate. Lane 5-8: Mdivi-1 treated TauTg mouse brain homogenate. Lane 9-10: Low dose Mdivi-1 treated TauTg mouse brain homogenate (0.01mg/month). Insufficient samples in low Mdivi-1 treatment group for statistical analysis. Lane 12: Ladder. Lane 13-14: WT samples for reference. Biological replicates.

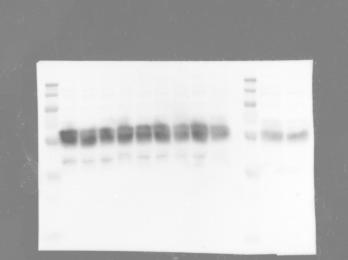

Mdivi-1

Veh

250 🡪

100🡪

55🡪

35🡪

Fig. S6. Raw image of representative western blot data shown in Fig 5D (2^nd^ row). Entire blot probed for pTau (S199/202) (50-64 kDa). Lane 1: 35 – 250kDa ladder. Lane 2-4: Vehicle treated TauTg mouse brain homogenate. Lane 5-8: Mdivi-1 treated TauTg mouse brain homogenate. Lane 9-10: Low dose Mdivi-1 treated TauTg mouse brain homogenate (0.01mg/month). Insufficient samples in low Mdivi-1 treatment group for statistical analysis. Lane 12: Ladder. Lane 13-14: WT samples for reference. Biological replicates.

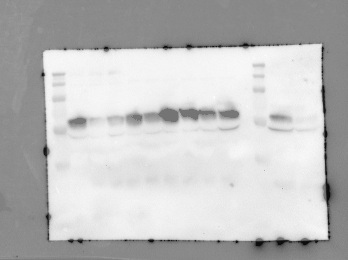

Veh

Mdivi-1

250 🡪

100🡪

55🡪

35🡪

Fig. S7. Raw image of representative western blot data shown in Fig 5D (3rd row). Entire blot stripped and reprobed for AT8 (64 kDa). Lane 1: 35 – 250kDa ladder. Lane 2-4: Vehicle treated TauTg mouse brain homogenate. Lane 5-8: Mdivi-1 treated TauTg mouse brain homogenate. Lane 9-10: Low dose Mdivi-1 treated TauTg mouse brain homogenate (0.01mg/month). Insufficient samples in low Mdivi-1 treatment group for statistical analysis. Lane 12: Ladder. Lane 13-14: WT samples for reference. Biological replicates.

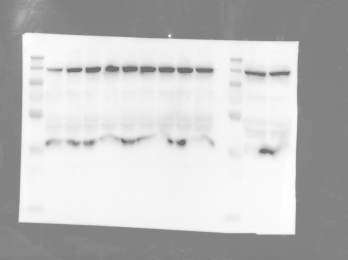

250 🡪

100🡪

55🡪

35🡪

Veh

Mdivi-1

Fig. S8. Raw image of representative western blot data shown in Fig 5D (4^th^ row). Entire blot stripped and reprobed for vinculin (124 kDa). Lane 1: 35 – 250kDa ladder. Lane 2-4: Vehicle treated TauTg mouse brain homogenate. Lane 5-8: Mdivi-1 treated TauTg mouse brain homogenate. Lane 9-10: Low dose Mdivi-1 treated TauTg mouse brain homogenate (0.01mg/month). Insufficient samples in low Mdivi-1 treatment group for statistical analysis. Lane 12: Ladder. Lane 13-14: WT samples for reference. Biological replicates.
